# A phenotypic screening platform utilising human spermatozoa identifies new compounds with contraceptive activity

**DOI:** 10.1101/808105

**Authors:** Franz Gruber, Christopher L.R. Barratt, Paul D. Andrews

## Abstract

There is an urgent need to develop new methods for male contraception, however a major barrier to drug discovery has been the lack of validated targets and the absence of an effective high-throughput phenotypic screening system. To address this deficit, we developed a fully-automated robotic screening platform that provided quantitative evaluation of compound activity against two key attributes of human sperm function: motility and acrosome reaction. In order to accelerate contraceptive development, we screened the comprehensive collection of 12,000 molecules that make up the ReFRAME repurposing library, comprising nearly all the small molecules that have been approved or have undergone clinical development, or have significant preclinical profiling. We identified several compounds that potently inhibit motility representing either novel drug candidates or routes to target identification. This platform will now allow for major drug discovery programmes that address the critical gap in the contraceptive portfolio as well as uncover novel human sperm biology.

## Introduction

An effective and comprehensive family planning strategy is fundamental to delivery on the United Nations 17 Sustainable Development Goals (Starbird *et al.*, 2016; Goodkind *et al.*, 2018), however the current contraceptive portfolio is suboptimal. For example, it is estimated that more than 214 million women in developing countries have an unmet need for contraception which results in 89 million unintended pregnancies and 48 million abortions every year, with substantial impacts on maternal and child health and poverty levels (Guttmacher Institute, 2017). To achieve the UN targets it is critical that involvement of men be considered of equal importance as women (Hardee *et al*., 2016). However, it is noticeable that no effective, reversible and widely-available form of contraception has been developed for men since the condom and as such the burden falls largely on the woman. The development of drugs that can be used in the male would address a critical gap in the contraceptive portfolio.

Progress in developing new male contraceptives has been slow for a plethora of reasons. Despite decades of research, safe and effective hormonal approaches have yet to be realised and viable alternatives remain elusive (Page and Amory, 2018). Developing a drug for any purpose is difficult but one that blocks sperm function is a particular challenge as spermatozoa don’t divide, transcribe or translate, have specialized organelles, are highly motile and continually produced in large numbers. Moreover, as the production of fertilization competent cells is necessarily different between species, it is essential to use a human system early in the drug discovery cascade to increase the chances of translational success.

Though some potential sperm-specific contraceptive drug targets have been identified (Lishko and Mannowetz, 2018), our relatively poor understanding of the spermatozoon makes target-agnostic strategies more attractive (Barratt *et al*, 2017). Phenotypic drug discovery is undergoing a renaissance in a number of therapeutic areas because it can uncover novel biology in an unbiased way. Retrospective analysis of drug approvals shows the approach to be more successful in first-in-class medicine discovery than previously appreciated (Swinney and Anthony, 2011; Moffat *et al*., 2017). For male contraceptive drug discovery, a limiting barrier has been the absence of a scalable drug screening system using spermatozoa.

To address this unmet need we developed the first high-throughput phenotypic sperm screening platform where an image-based kinetic analysis module, tracking movement, is coupled to a flow cytometry module that detects the acrosome reaction (AR), using a fully-automated robotic system. The bespoke computational pipeline quantitatively evaluates compound activity against both attributes thus achieving unprecedented throughput. The platform was used to identifying inhibitors from a unique 12,000 molecule ReFRAME (Repurposing, Focused Rescue, and Accelerated Medchem) library, which represents the most comprehensive collection of drugs available, as it contains nearly all the small molecules that have achieved regulatory approval and others that have undergone varying degrees of clinical development or have had significant preclinical profiling (Janes *et al.*, 2018). This advance opens up the possibility of accelerated male contraceptive development by allowing drug repurposing, target identification as well as screening of chemical diversity libraries for novel medicinal chemistry start points.

## Results

### Phenotypic Assay Development

Our automated platform (Figure 1A) allows for the identification of relatively fast-acting compounds with a rapid plate-processing time (~75 minutes/ 384-well plate), where the motility assay runs for ~30 minutes followed directly by the AR assay (~ 45 minutes). Automated cell tracking (Figure 1B & Supplementary Figure 1A) using time-lapse images allowed kinetic parameter quantification on >200 cells/well (Figure 1C). The kinetics show good agreement with clinical systems (Supplementary Figure 1B) and a comparable distribution was observed between experimental days, plates and positions (Supplementary Figure 1.1 A) with any small differences between days stemming from donor pool variance. Importantly, spermatozoa were found to be tolerant to DMSO (~4%), far above typical screening concentration of 0.1% (Supplementary Figure 1C). To determine the optimal screening batch size spermatozoa were dispensed every 30 minutes for 150 minutes observing only a small decrease in motility (~10%) after 4 plates were screened (Supplementary Figure 1D). A flow cytometry-based assay measuring AR (Figure 1D) was run directly afterwards (Figure 1E, F & Supplementary Figure 1E).

**Figure 1.**
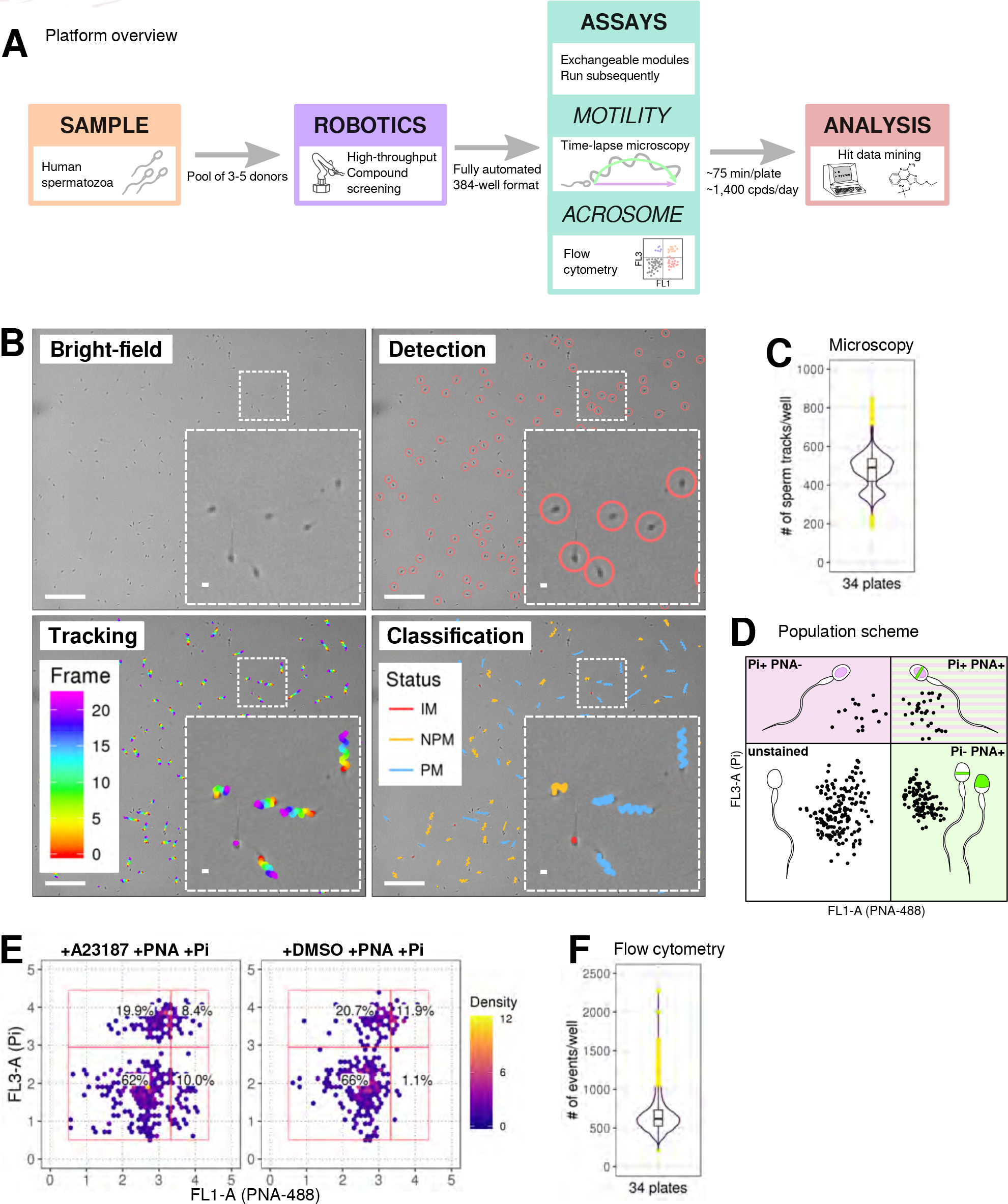
Phenotypic Assay workflows. (**A**) Graphical summary of modular screening workflow where motility measurement is followed by acrosome reaction (AR) measurement allowing a screening throughput of >1400 compounds per donor pool (**B**) Steps in imaging and analysis: human sperm are recorded with brightfield illumination (first panel) then sperm heads detected using a particle tracking algorithm (second panel) which are then tracked across the timelapse series of images (third panel) and subsequently classified (fourth panel). Each panel contains a zoomed-in subsection of the field. Colour coding for tracking distance and kinetic classification is shown in the panel insets: *Tracking* panel - rainbow gradient (showing progression over time); *Classification* panel - red for immotile (IM), yellow for non-progressively motile (NPM) and blue for progressively motile (PM). Scale bars: 100 μm (main images), 5 μm (insets). (**C**) Sperm counts per well after microscopy and detection shown in a combined violin/box plot. Colors: purple violin outline (probability density of values), yellow dots (outliers of boxplot). (**D**) Graphical summary of the expected populations determined by flow cytometry based on distribution of cells measured with FL3-A (Pi) vertical axis and FL1-A (PNA-488) horizontal axis: dead cells (upper left, Pi+ PNA−); dead and acrosome-reacted (upper right, Pi+ PNA+); unstained/live/non-reacted (lower left) and live acrosome-reacted (lower right, Pi-PNA+). (**E**) Example flow cytometry data comparing sperm treated with the Ca^2+^ ionophore (A23187) which induces AR (left panel) with sperm from DMSO-treated well (right panel). Colors indicate event density. (**F**) Combined violin/box plot data showing flow cytometry event counts per well. Colors and label as in (**C**).

### Screening of the ReFRAME library, confirmation of hits by dose response

The ReFRAME library (11,968 compounds) was supplied in “assay-ready” imaging plates and was solubilised prior to addition of sperm. A total of 63 compounds decreased motility (Figure 2 A, B with examples in Figure 2C, E and supplementary movies 1 & 2). High assay quality was achieved throughout with a Z’-factor between 0.4-0.8 (Figure 2D). 14 compounds were selected as AR^+^ hits (Figure 3 A with examples in B, C). Motility hits were confirmed with resupplied material in dose-response experiments. There was a dose-dependent decrease in motility for 29 compounds, with EC_50_ values as low as 0.05 μM and effect sizes ranging from 15-100% (Figure 2 F, G; Supplementary Figure 2, Supplementary Table 1). Amongst the confirmed hits were the aldehyde dehydrogenase inhibitor, Disulfiram (70% maximum reduction at 10 μM) and a putative platelet aggregation inhibitor, KF-4939 (100% max. reduction; EC_50_=0.49μM), and a range of other compounds having very modest effect and/or showing low potency (See Supplementary Figure 2, Supplementary Table 1). In the AR screen, 9 of the resupplied compounds had a dose-dependent effect with EC_50_ values as low as 0.4 μM (See Supplementary Figure 3, Supplementary Table 2). However, following the orthogonal assay triaging none of the AR hits were found to be true positives (see examples in Supplementary Figure 3.1 D).

**Figure 2.**
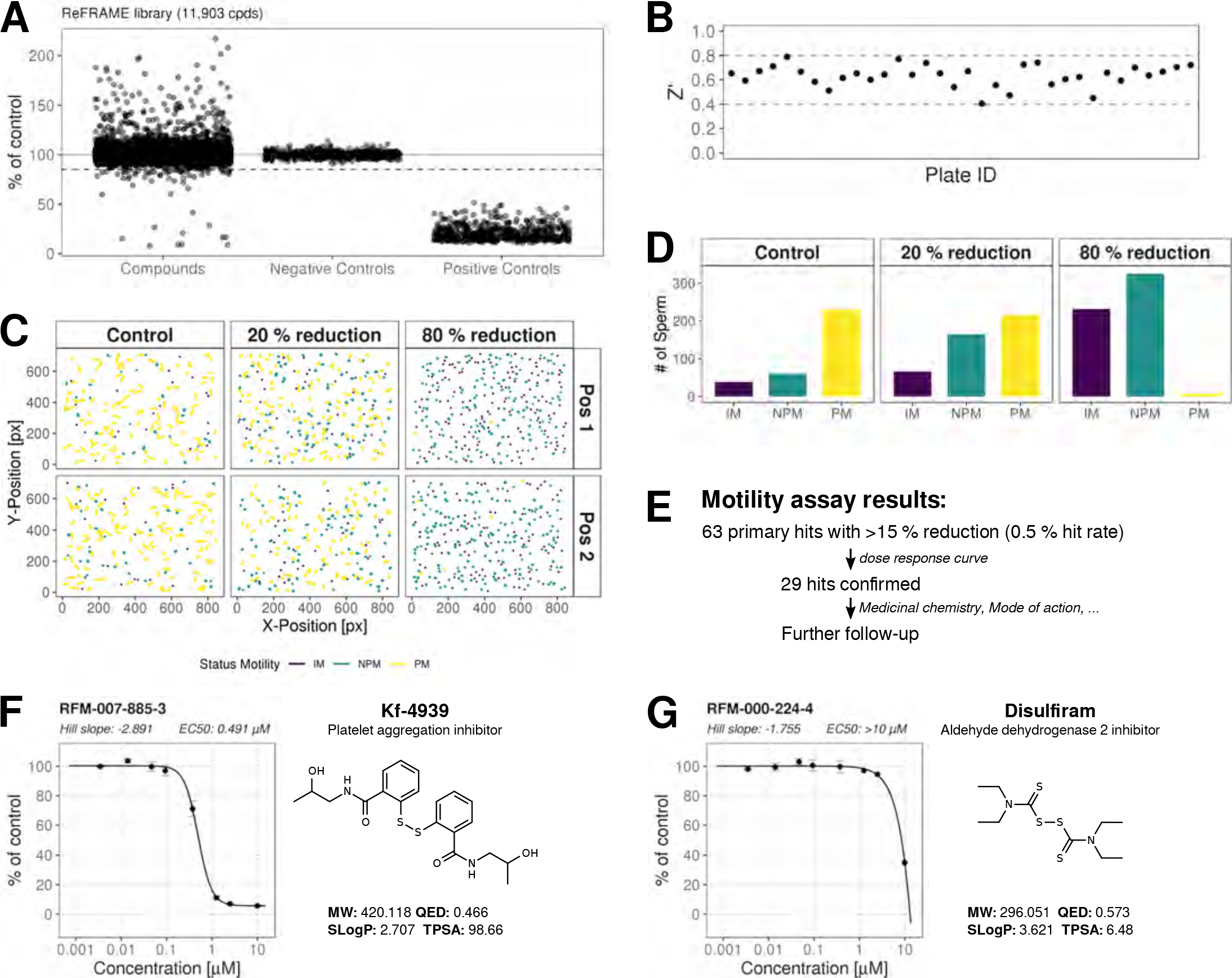
ReFRAME Library Screening: Motility Assay Results. (**A**) Primary screening results of the motility assay. Each dot represents a well (either compound or control well) showing % of DMSO control (VCL). Positive controls (Pristimerin), negative controls (DMSO) and individual compound datapoints are shown. The dashed line showing the 15% of control (reduction in VCL) – the cut-off for primary hit selection. Total number of compounds = 11,903, excluding wells with auto-focus errors and with “sticky” compounds which have been excluded from analysis. See Figure_2_Source_Data.H5 along with Figure_2_Code.R. (**B**) Assay robustness metrics: Z'-factor values for all screening plates is shown. Dashed lines indicate min/max Z’ values. (**C**) Tracking data visualizations of 3 example wells showing sperm tracks of both imaging positions (Position 1 [Pos. 1] and Position 2 [Pos. 2] respectively) within the wells. A DMSO control well (left panels “Control”) shows a large number of progressively motile (PM) sperm (yellow) with few non-progressively motile (NPM) sperm (green and very few immotile (IM) sperm (purple) – this is in contrast to the shorter tracks and higher levels of NPM and IM in the middle panel (“20% reduction”) for a compound that shows 20% inhibition of motility (i.e. 80% of control) and the right hand panel (“80% reduction”) for a compound showing 80% inhibition (i.e. 20% of control), showing almost all cells are in the IM and NPM classes. (**D**) Histogram of sperm tracks quantification of the data show in (**C**). **(E)** Summary of motility assay hit rate (0.5%) and reconfirmation rate (0.24%). (**F-G**) Dose response confirmation of two hits. 8-point 3-fold dilution curves are shown with 10 M as the highest concentration. two data points per concentration (n=2, data point is mean ± SD). Each curve is a 4-parameter logistic fit. Each plot shows estimated values Hill Slope and EC_50_. The chemical structure of the hit compound is shown as well as some annotation and physicochemical properties. Physicochemical properties were calculated using RDKit, Python and KNIME: SlogP = partition coefficient (Wildman and Crippen, 1999); TPSA is the Topological Polar Surface Area (Ertl *et al*., 2000); MW is the exact Molecular weight; QED= Quantitative Estimate of Drug-likeness (Bickerton *et al* 2012).

**Figure 3.**
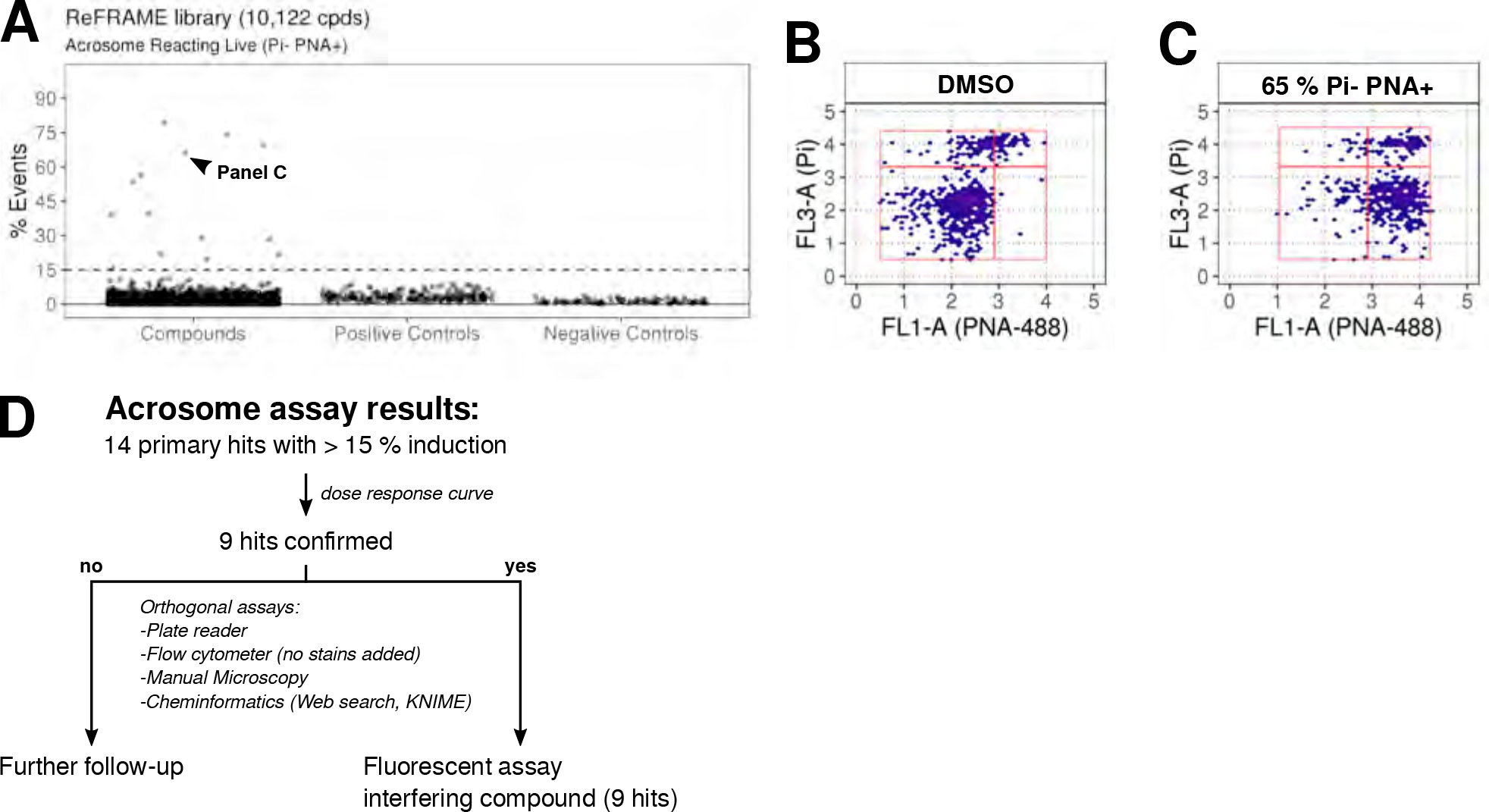
ReFRAME Library Screening: AR Assay Results. (**A**) Results of primary screening of the library using the acrosome assay (live cells, acrosome reacting, Pi− PNA+ population). Each dot represents a well (either compound or control well). Shown is % Events (number of events in Pi− PNA+ gate relative to total events in the sampled well). Datapoints for compounds, negative controls (DMSO) and positive controls (A23187) are shown. Black dashed line = 15% threshold for primary hits selection. See Figure_3_Source_Data.csv along with Figure_3_Code.R. (**B-C**) Data from two example wells: a DMSO well (left panel) and a well with 65% Pi− PNA+ population (right panel). **(D)** Summary of acrosome assay results before and after triage.

## Discussion

A major barrier in the search for new male contraceptives has been the lack of an effective high throughput phenotypic screening system. This study presents the first platform based on imaging and flow cytometry that measures two fundamental aspects of sperm behaviour – motility and AR, allowing for the rapid screening of compound collections.

A significant challenge was to create a system for the tracking of these highly motile cells that was fully automated and scalable to allow throughput at sufficient speed. Moreover, concomitant assessment of a second functional attribute, AR, was developed in order to maximise use of the biological material. Assessment of motility required novel implementation of tracking algorithms. Importantly, the kinetic outputs were equivalent to those from the “industry-standard” CASA, which is very low throughput and unsuitable for screening more than a few compounds at one time. Significant workflow optimisation resulted in good assay robustness and a high hit confirmation rate following the primary screen. For AR detection, a flow cytometry approach was taken based on established methods (Mortimer *et al.*, 1987). However, the presence of assay-interfering compounds in screening libraries (potential fluorescent compounds including DNA intercalators) necessitated further triaging steps (see Materials and Methods). This combination of triaging approaches meant that, in this screen, no AR compounds were suitable for progression. Overall, although this was a complex biological assay, it achieved acceptable throughput with the potential to increase batch size for larger screening campaigns.

The advantage of using the ReFRAME collection was the potential to repurpose drugs that are already approved for other indications or are at earlier stages of clinical progression. The associated compound annotations and safety data offers the potential to help accelerate the development of an effective male contraceptive. In the motility assay the hit rate of ~0.2% is typical for a cell-based assay, however surprisingly a relatively high number of hits were not deemed suitable for compound progression. Examples included potentially toxic mercury-containing compounds (phenylmercuric borate and mercufenol chloride); antiseptics and antibiotics including tyrothricin, or compounds with impacts on fundamental biology e.g. the microtubule stabiliser and chemotherapeutic, docetaxel. However, several hits are potentially of interest. One is Disulfiram (70% maximum reduction at 10 μM), a drug that is well tolerated, has long been used for treating alcohol dependency and has previously been shown to inhibit sperm motility (Lal *et al.*, 2016). Disulfiram is unlikely to be developable as a suitable contraceptive drug due to its side-effects when alcohol is consumed but has potential to facilitate mode-of-action studies. A recent report identified 3-Phosphoglycerate dehydrogenase (PHGDH) as a target for disulfiram (Spillier *et al.*, 2019) although the relevance to sperm function is unclear. The identification of KF-4939, a putative platelet aggregation factor (PAF) inhibitor, as a potent hit (100% max. reduction EC_50_ = 0.5 μM) may also be of interest since PAF has been used to improve sperm function in male infertility treatments (Roudebush *et al.* 2004). It is noteworthy that like disulfiram, KF-4939 also contains a disulphide linkage indicating that its biological effect maybe mediated by modifying sulfhydryls on proteins (Yamada *et al.*, 1985). Further thiol-containing compounds, ethanedithiol (98% max reduction; EC_50_=2.5μM) and the anti-fungal benzothiazole, Ticlatone (96% max. reduction EC_50_ = 4.4 μM) were also potent hits, suggesting free cysteines might play a critical role in motility.

Moreover, the Toll-like receptor 7/8 ligand (Resquimod; R848) recently shown to preferentially reduce the motility of X chromosome-bearing mouse sperm by suppressing ATP production (Umehara *et al*, 2019) was a hit in our screen, demonstrating that this phenotypic screening approach has the potential to uncover new aspects of sperm biology.

In summary, our high-throughput phenotypic platform allows for the screening of bioactives including natural products, focussed small molecule libraries and chemical diversity sets, which can provide novel start-points for male contraceptive drug development.

## Material and Methods

### Ethical approval

Written consent was obtained from each donor in accordance with the Human Fertilization and Embryology Authority (HFEA) Code of Practice (version 8) under local ethical approval (13/ES/0091) from the Tayside Committee of Medical Research Ethics B.

### Development of methods for Motility and Acrosome Reaction (AR)

A pre-existing automated cell-based phenotypic screening platform was adapted that utilises a Yokogawa CV7000 Cell Voyager high-throughput microscope able to image 384 multiwell plates under full environmental control at high speed. For high throughput screening with human spermatozoa to be effective and yield meaningful hits, the system needed to replicate, as close as possible, the manual workflow currently used in an andrology lab, so that the effect of large numbers of compounds on viable human sperm could be performed. In order to achieve this goal, we investigated a number of factors that could impact the quality of the data such as: plate type; concentration and number of spermatozoa per well; logistics of spermatozoa preparation and handling; dispense speed for getting spermatozoa into 384 well plates; timing of compound addition; temperature control; image acquisition parameters; algorithm choice for motility assessment and data management and, evaluating the overall speed of data analysis. A flow cytometry-based assay and data analysis workflow was developed to measure acrosomal status. The goal was to be able to screen a 384-well plate in both assays consecutively within 90 minutes of sperm dispensing using a fully automatic robotic system. Data was analysed for inter- and intra-plate variation (both technical and biological replicates) and signal-to-background and variance measured to demonstrate the assay was reproducible and robust. The assay was shown to be scalable and all the workflows optimal. Once established, the system was validated by screening the ReFRAME library (Janes *et al.*, 2018) and hits confirmed by dose-response analysis.

### Sperm handling

Control semen samples were obtained from volunteer donors with normal sperm concentration, motility and semen characteristics (WHO, 2010) and no known fertility problems. Samples were obtained by masturbation after 48–72 h of sexual abstinence. After ~30 min of liquefaction at 37°C, cells were prepared using a discontinuous density gradient procedure (DGC). Semen was loaded on top of a 40%-80% suspension of Percoll (Sigma Aldrich, UK) diluted with non-capacitation medium using Minimal Essential Medium Eagle, supplemented with HEPES, Sodium lactate and Sodium Pyruvate to achieve a similar buffer as described previously (Tardif *et al.*, 2014). We routinely pooled samples from 3-5 donors for use in each screening batch run in order to reduce donor-to-donor variability. Samples were obtained and analysed in line with suggested guidance for human semen studies where appropriate (Björndahl *et al.*, 2016).

### Motility assay

Sperm cells were incubated for 3 hours at 37°C after DGC, transferred to the robotic platform (HighRes Biosolutions Inc.) and maintained at 37°C with gentle stirring (100 rpm) in a water bath on top of a magnetic stirrer. Approximately 10,000 spermatozoa were dispensed per well using a MultiDrop Combi (ThermoFisher) into pre-warmed assay-ready plates. Spermatozoa were incubated with the compounds for 10 mins (thus favouring fast acting compounds) at 37°C in the CV7000 microscope prior to the commencement of imaging.

### Time-lapse Imaging

A CV7000 Cell Voyager high-content imaging system (Yokogawa) was used as it allowed full environmental control, sufficient contrast using simple brightfield optics and a fast acquisition rate (up to 45 frames per second). Using a 20× lens with bright field illumination (0.11 ms exposure, 3% lamp power), time-lapse image series were acquired (24 frame in 0.5 sec) at two positions per well (with a 400 μm gap between positions to eliminate double counting of sperm). Image acquisition across a 384-well plate with such settings takes ~17 minutes.

### Sperm Tracking

A Python implementation of the particle tracking algorithm originally developed by others (Crocker and Grier, 1996) was employed (Trackpy v0.4.1 Allan *et al.*, 2018). The algorithm determines the position of every sperm head in each frame and links the position over time creating tracks. We optimized the algorithm parameters for our imaging data to exclude sperm cross-tracks and avoid detecting non-sperm particles. The image data (~17,000 files amounting to 21GB per plate) can be processed within 30 minutes using a standard desktop PC (Intel® Core^™^ i5-6600 CPU, 3.3 GHz, 8 Gb RAM) and can also be parallelized on a compute cluster with minimal effort.

### Acrosome assay

A flow cytometry-based assay was developed employing an iQue Screener (Sartorius) with Alexa488-conjugated peanut agglutinin (PNA) (Mortimer et al., 1987) and general sperm viability (using propidium iodide, Pi). This assay was run directly after motility assessment. Controls (DMSO and the Ca^2+^ ionophore A23187) were added using an acoustic dispenser (Echo 555, Labcyte Inc.) and incubated for 10 minutes at 37°C in the SteriStore (HighRes Biosolutions Inc.). PNA488 (ThermoFisher Scientific, Cat. No. L21409, stored as 1 mg/mL stock) was then added to achieve a 1:1000 final dilution and propidium iodide (ThermoFisher Scientific, Live/Dead Sperm Viability kit, Cat. No. L7011, stored as 2.4 mM solution) was added to achieve a 1:2000 final dilution using a liquid handler (Tempest, Formulatrix). After addition of dyes, plates were incubated for 10 minutes at 37°C in the SteriStore before sampling using an iQue Screener (2 sec sip time per well with pump setting of 45 rpm). This resulted in a sample processing time of about 25 minutes per 384-well plate. Cells were categorised as either: unstained cells; “dead” cells (Pi+ PNA−); Acrosome-reacted but “dead” cells (Pi− PNA+); or Acrosome-reacted “live” cells (Pi-PNA+). This method is capable of detecting a shift of populations (Pi+ and PNA+) upon induction of AR using A23187. The AR assay was performed as an agonist screen (scoring for induction of AR in live cells compared to DMSO controls) with the expectation that compounds could be found that induce AR beyond levels of induction with A23187.

### Controls and QC criteria

For the motility assay DMSO was used as a vehicle control (negative control) and Pristimerin (Merck, Cat. No. 530070, stored as 10 mM stock in DMSO; final concentration of 20 μM) as a positive control. For the acrosome assay we added a calcium ionophore (A23187, Sigma-Aldrich, Cat. No. C7522, stored as 10 mM stock in DMSO, used at a final concentration of 10 μM) as a positive control and DMSO as a negative control. We calculate a Z’-factor (Z’; Zhang, 1999) for each plate using positive and negative controls for the motility assay and observed Z’-factors ranging between 0.4-0.8. In addition we performed a visual check of heatmaps for every plate to detect edge effects.

### Data analysis and normalization

**Motility:** custom R scripts were written to calculate standard sperm kinetic parameters (Mortimer *et al.*, 2015). Those parameters allow classification of sperm into standard WHO classes: progressively motile (PM) (where VAP > 25 um/sec AND STR > 80%); non-progressively motile (NPM) (where VAP > 5 um/sec OR VSL > 11 um/sec), and immotile (IM). In addition to calculating kinetic parameters, we established a workflow to generate movies of time-lapse videos with overlapping sperm tracks using R and FFMPEG (FFmpeg Developers. Available from http://ffmpeg.org). We expressed results as % of control VCL. This was defined as VCL_median (cpd)/VCL_median(DMSO) * 100. Hit selection criteria was 15% reduction of VCL. VCL was chosen as the main kinetic measurement as it had an acceptable Z’ value and is independent of path averaging. Wells with autofocus errors or compounds which did not dissolve properly (resulting in sperm cells being stuck in one location but moving normally in the rest of the well) were excluded from analysis.

**Acrosome:** iQue Screener Data was exported as FCS format and processed using the following *Bioconductor* packages for analysing flow cytometry data: flowCore, flowDensity, flowWorkspace, ggcyto (see https://bioconductor.org). These packages allow handling flow cytometry data as objects, which can then be compensated, gated and visualized efficiently. In addition we use HDF5 format (www.hdfgroup.org) for storing processed and averaged data. Percent events for each population (Pi+ PNA+, Pi-PNA+, Pi+ PNA- and Pi-PNA−), normalized to total well events, were calculated. Hits were defined as compounds which induce AR beyond 15% (maximum level of induction achieved with positive control). Wells with irregularities (low in Pi+ or Pi+PNA+ population, or below 200 events) have been excluded from analysis.

### Compound Screening

We screened the ~12,000 compound ReFRAME (Repurposing, Focused Rescue, and Accelerated Medchem) library (Janes *et al.*, 2018; and www.reframedb.org) supplied by CALIBR at the Scripps Institute. This unique library consists of bioactives and approved drugs that have been assembled from the literature, drug databases and by patent mining. Compounds were spotted (12.5 nL, final assay concentration ~6 μM) into 384-well black-sided optical imaging plates (CellCarrier, PerkinElmer) at CALIBR and were shipped on dry ice to the screening centre in Dundee and stored at −20°C until required. Immediately prior to screening plates were thawed, controls were added using the acoustic dispenser (Echo 555, Labcyte), and 10 μl warm media added followed by shaking on a plate shaker for 10 seconds. Plates were then incubated for 30 min at 37°C to solubilise the compounds and prewarm the plates prior to the addition of live sperm.

All experiments were performed with pools of donors (3-5 per batch of 4 screening plates) and results normalised to DMSO-treated wells (16 wells per plate).

The ReFrame library has been screened for cytotox (using CellTiterGlo) in HEK293 and HepG2 cells (see www.reframedb.org) with some of the hit compounds displaying activity in one or both assays (Supplementary Figure 2), information which helps inform compound progression decisions.

### Dose response experiments

Assay-ready 384-well CellCarrier plates with 8-point curves (10 μM highest concentration) were resupplied by CALIBR in duplicate and processed as described above. Curves were fitted using the R package *dr4pl* (https://cran.r-project.org/web/packages/dr4pl/index.html) using a 4-parameter logistic fit option.

### Triaging of AR hits

Given the prevalence of a range of interfering compounds amongst the library hits, two approaches were used for triaging. These additional triaging steps, after the primary screen, allowed for the elimination of intrinsically fluorescent compounds and compounds fluorescing in the presence of biological material. Screening plates were excited at 488 nm and read at 520 nm and 670 nm in a M1000Pro multimode plate reader (Tecan) after flow cytometry. In addition, when performing dose-response curve follow up on resupply material a replicate set of compound-dosed cells were processed with no-dyes added. This identified the small number of compounds whose fluorescence is indistinguishable from a true positive. In addition, visual confirmation of the appropriate pattern of acrosome staining was performed by microscopy.

---

**Supplementary Figure 1.**
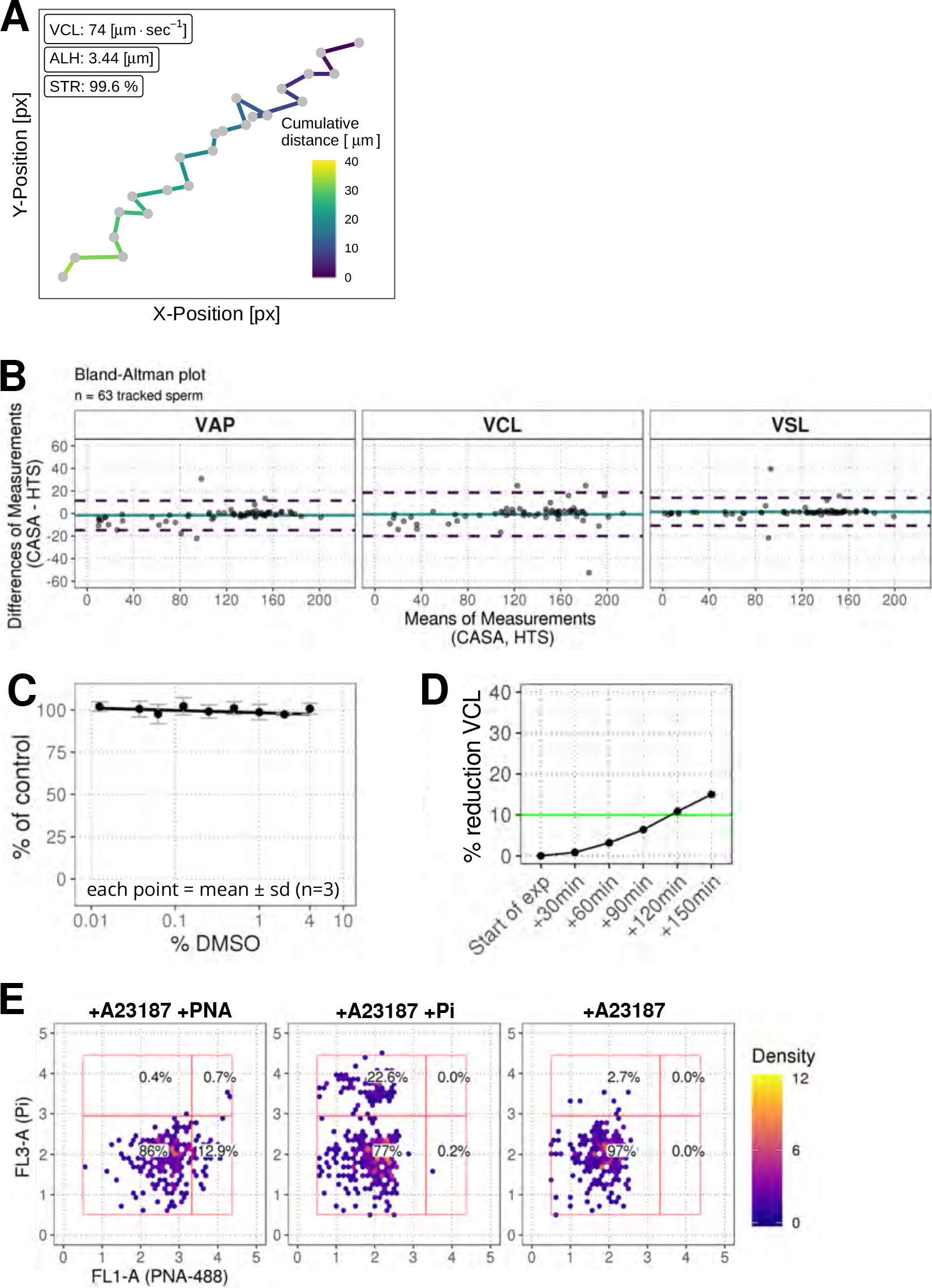
Further characterisation of phenotypic assays. (**A**) Example data for a single sperm track with *x* and *y* coordinates being used to calculate standard sperm kinetics. Color represents the cumulative distance travelled in microns. Acronyms: VCL (curvilinear velocity), ALH (amplitude of lateral head displacement; maximum value), STR (straightness ratio). (**B**) Comparison of data from the standard computer-assisted semen analysis (CASA) with the high-throughput system using a Bland-Altman plot of VAP, VCL, and VSL. Colors: turquoise line (mean of differences), dashed purple lines (limit of agreement, mean of differences +/− 1.96 * SD). (**C**) Effect of DMSO on sperm motility (VCL) relative to untreated wells. (**D**) Effect of pre-dispense incubation time on sperm motility (% reduction in VCL). Colors: green line (arbitrary 10% cut-off). (**E**) Plots of FL3-A vs FL1-A for the flow cytometry assay controls using acrosome specific PNA-488 dye (FL-1) and cell viability marker propidium iodine (Pi, FL-3) upon addition of A23187 Ca-ionophore. Panels from left to right: with ionophore and acrosome stain; with ionophore and propidium iodide only, with ionophore but unstained. Colors: red rectangles show population gates; colored hexagons show event density (bin size = 40).

**Supplementary Figure 1.1.**
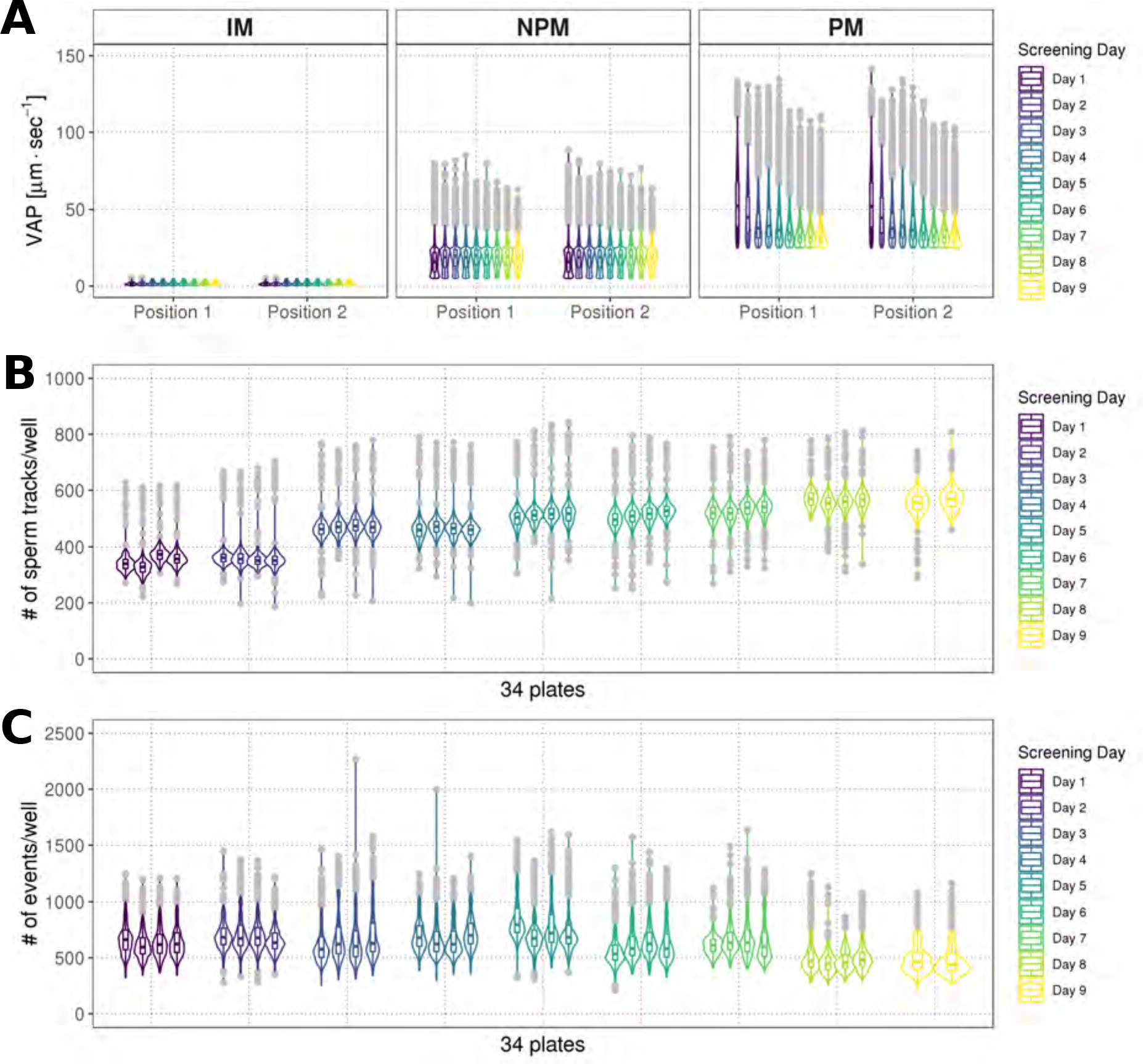
Screening consistency over time analysis. (**A**) Distribution of average path velocities (VAP) for sperm motility classes (IM = immotile; NPM = non-progressively motile; PM = progressively motile), using data from two positions within each well (Position 1, Position 2), represented as combined box/violin plot. Each box/violin plot is a summary of a screening day (n = 4, 384-well plates). Colors: each screening day is represented as a different color; grey = boxplot outliers. (**B**) Distribution of numbers of tracked sperm per well across screening days, represented as combined box/violin plot. Each box/violin plot is a summary of one screening plate. Colors as in (**A**). (**C**) Distribution of events identified in the flow cytometry assay, represented as combined box/violin plots. Each box/violin plot is a summary of a screening plate. Colors as in (**A**).

**Supplementary Figure 2.**
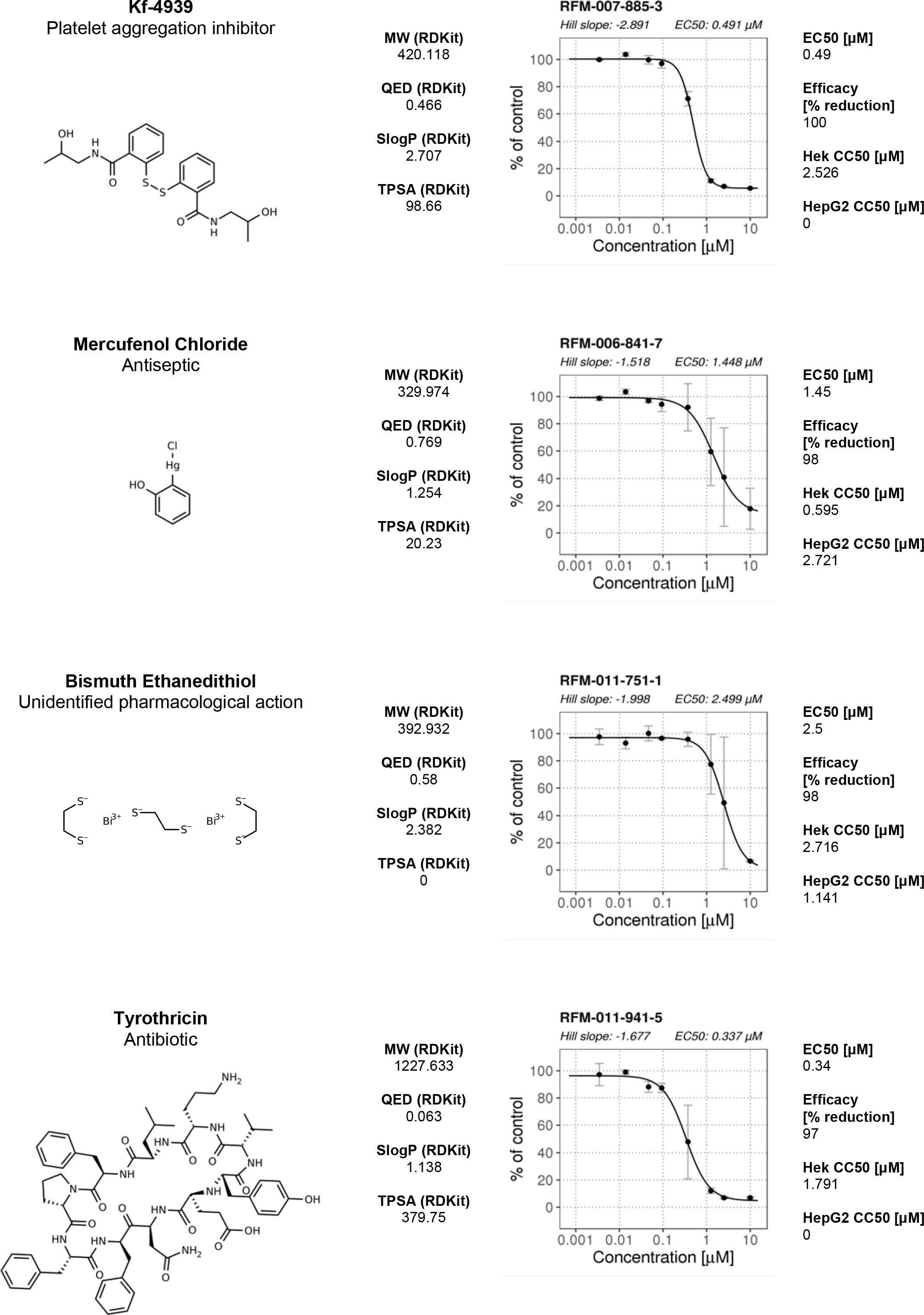
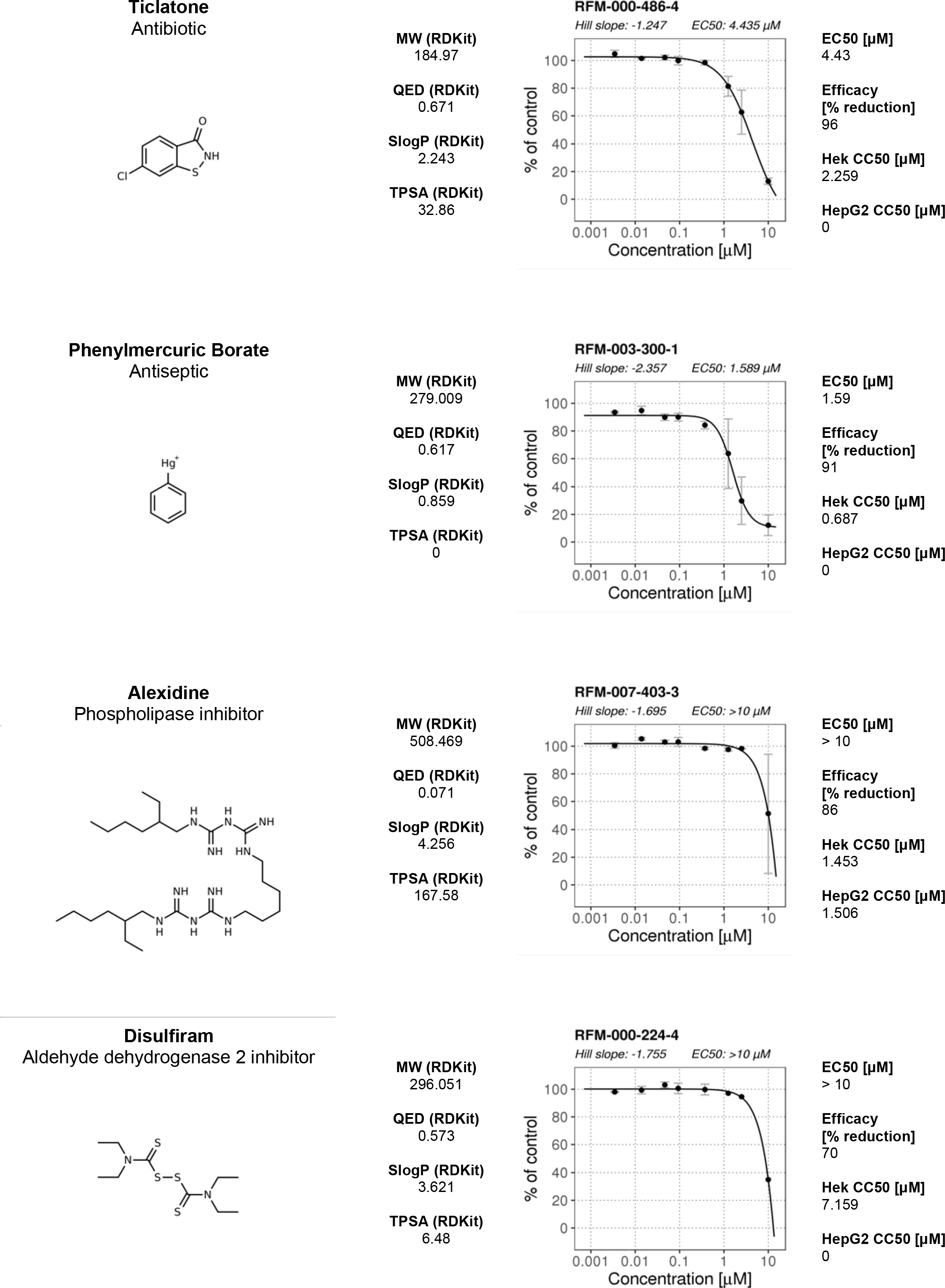
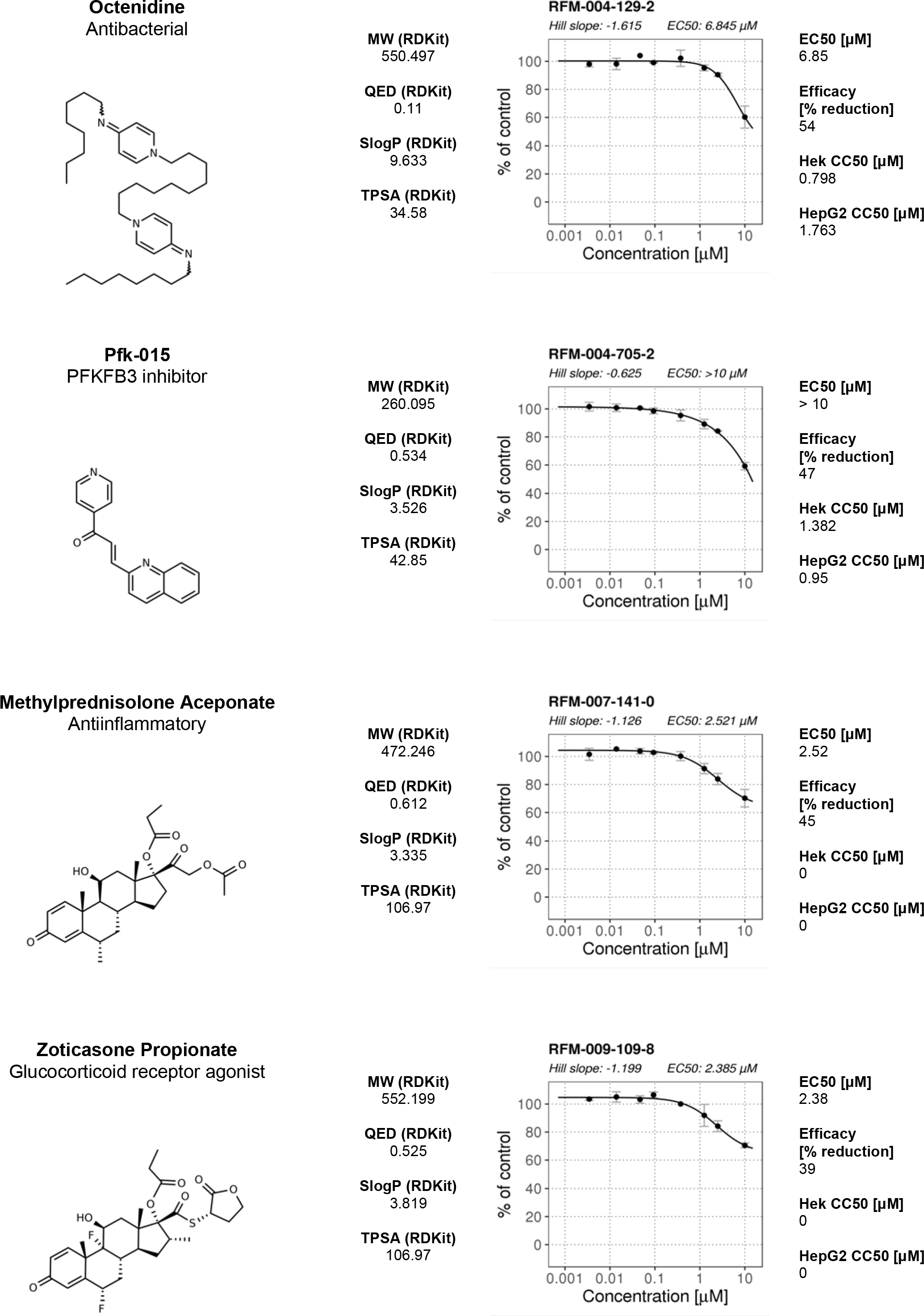
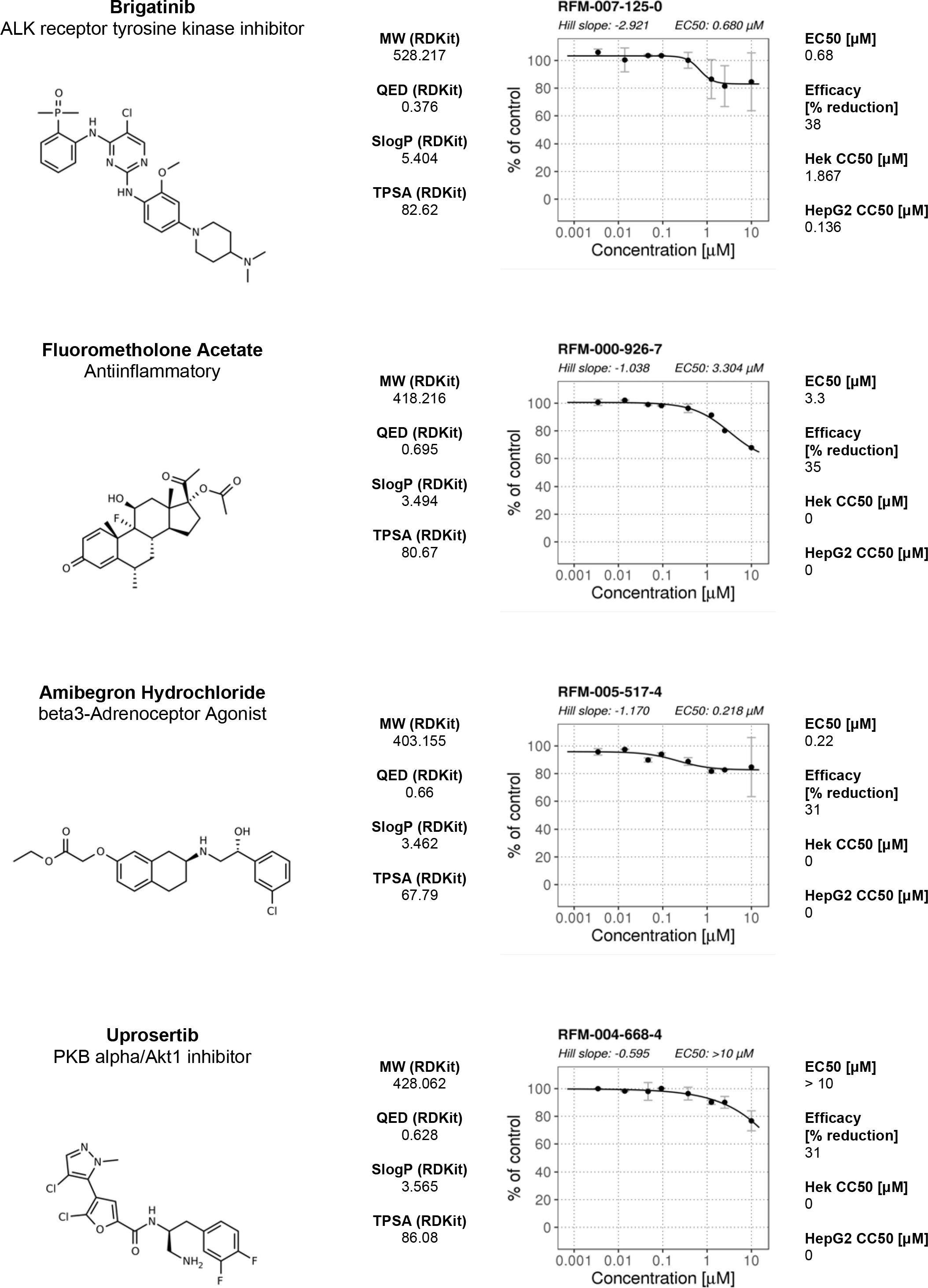
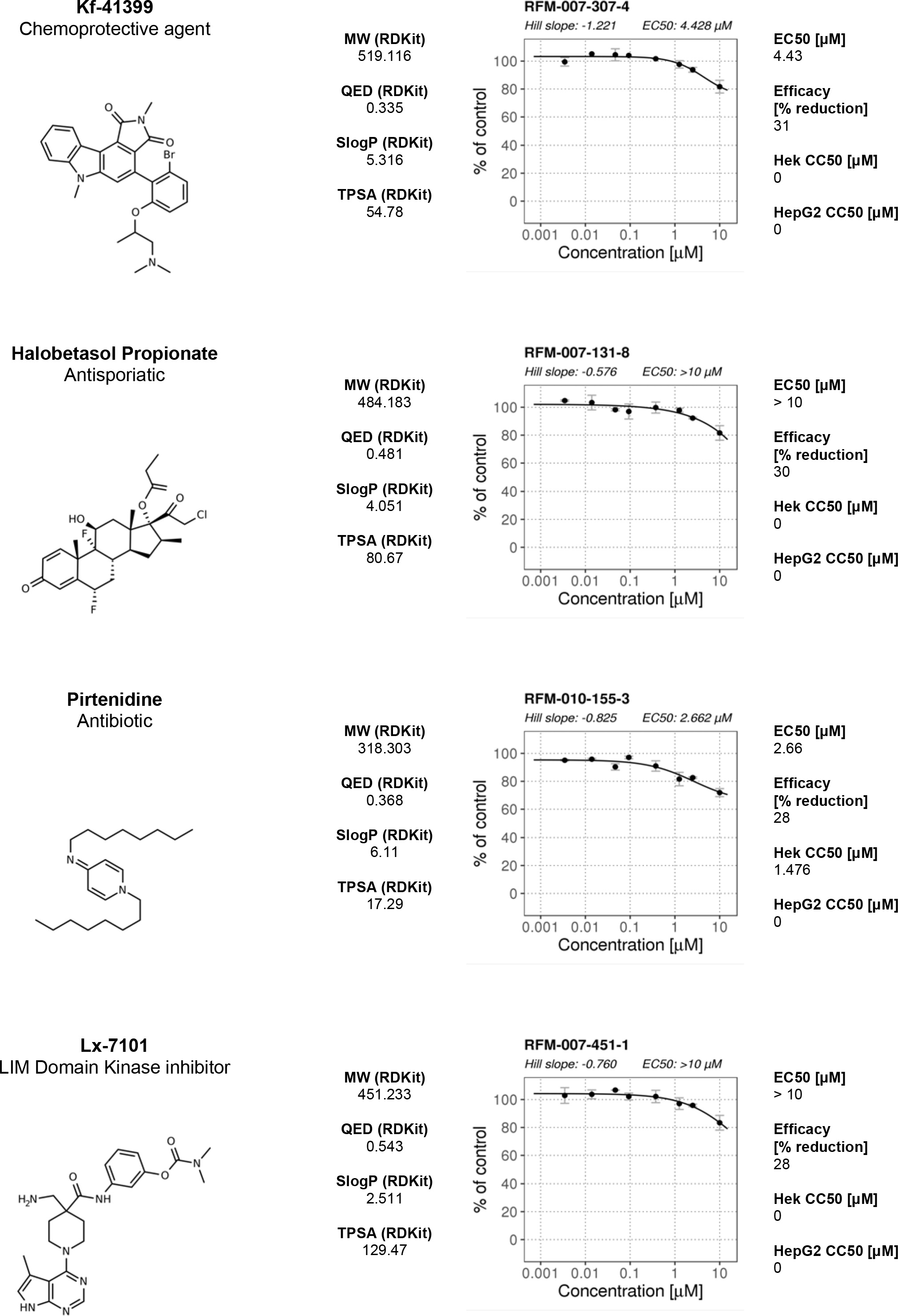
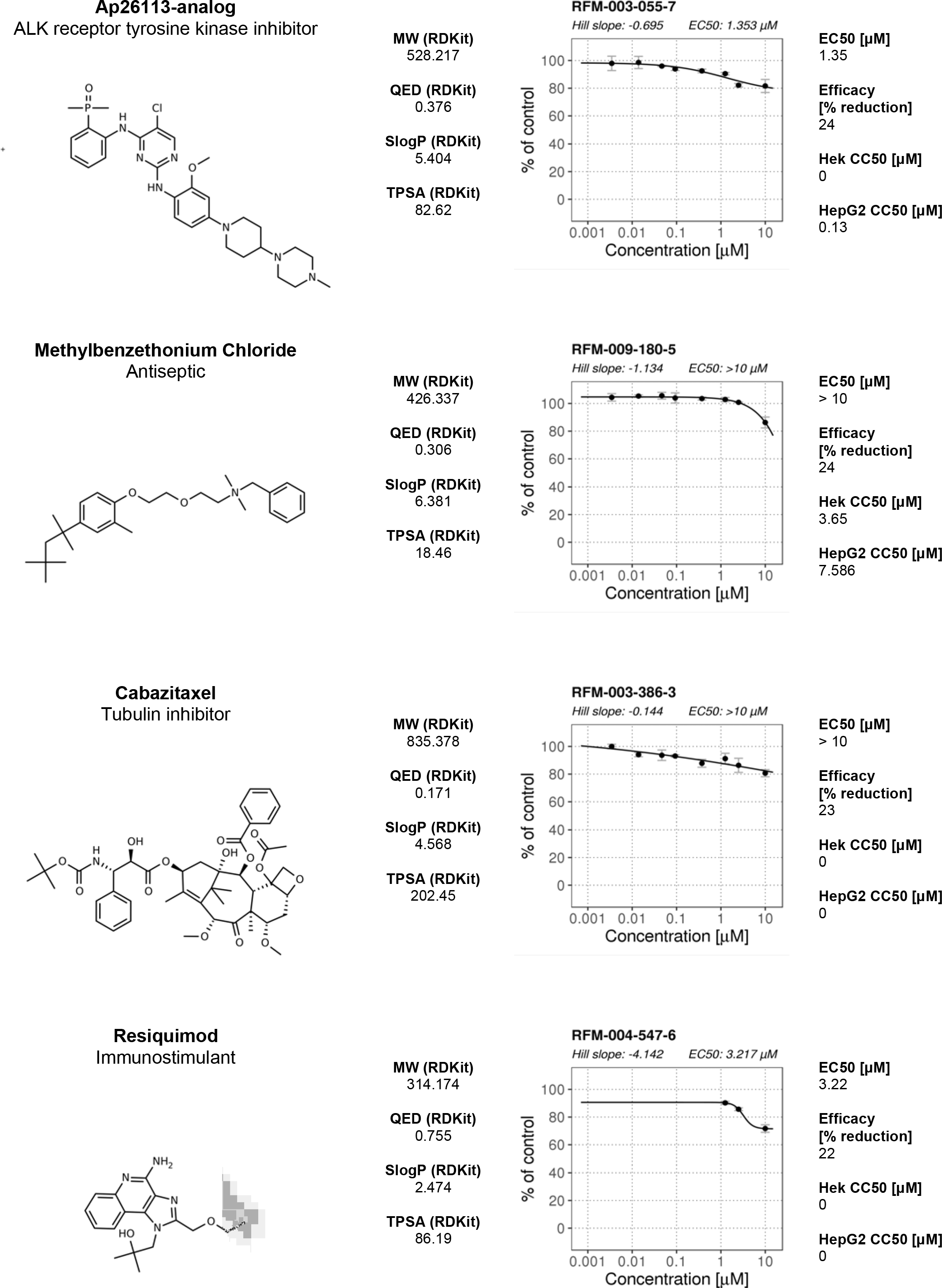
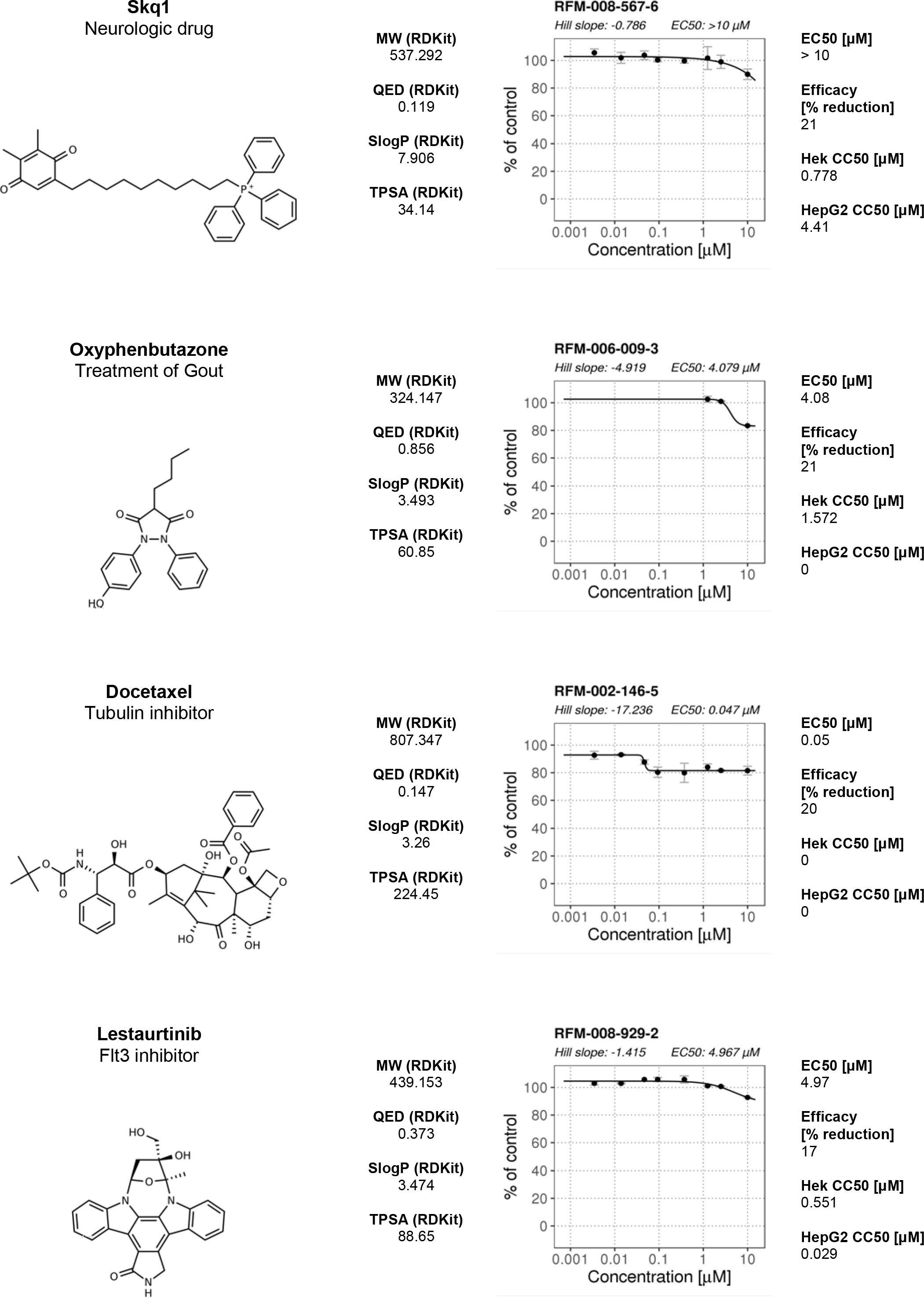
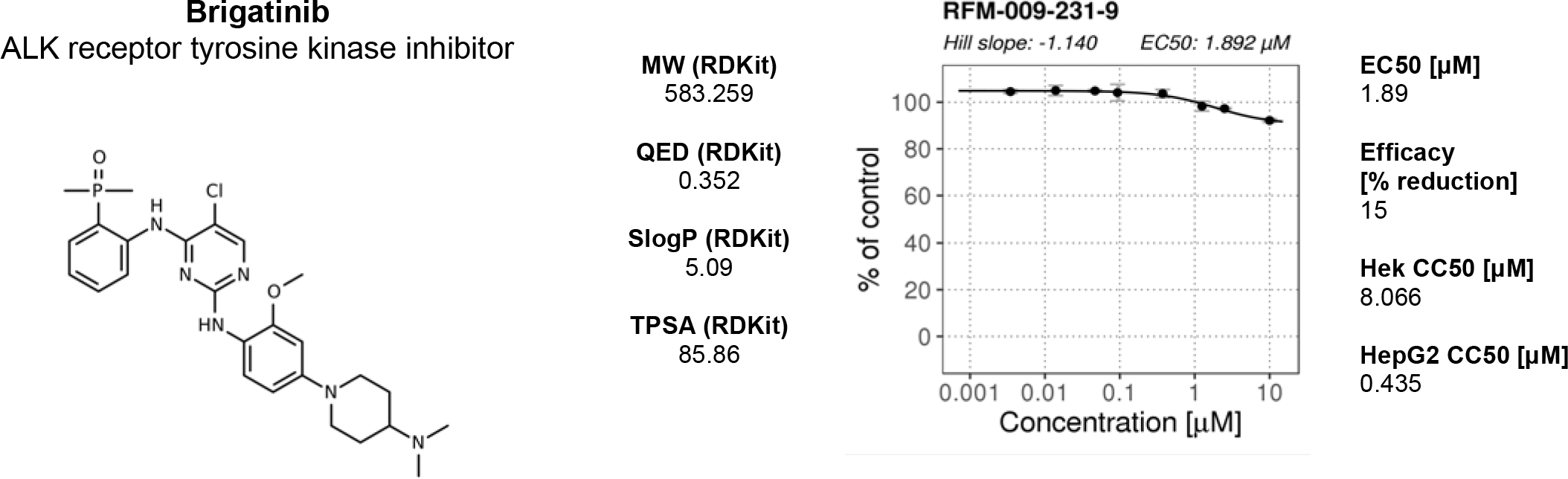
Dose response and additional data for primary motility hits. Compound names and nominal target/functional annotation shown on left along with chemical structure and physico-chemical properties. The 8-point dose response curves for each hit with estimated Hill slope, EC_50_ and Efficacy [% max reduction] are shown on the right. Two data points per concentration (n=2); data points are Mean ± SD. A 4-parameter logistic fit has been performed using R package: *dr4pl*. Calculated EC_50_ values and Efficacy (% max effect) in the motility assay are shown in the right hand section. CC_50_ values for HEK293 and HepG2 CellTiter-Glo cytotoxicity assays performed by CALIBR (www.reframedb.org) are shown in the right hand section. Note: 0 = inactive in cytotoxicity assay. The chemical structure of the hit compound is shown as well as some annotation and physicochemical properties (for nomenclature description see Figure 2). See Supplementary_Figure_2_Source_Data.csv along with Supplementary_Figure_2_Code.R.

**Supplementary Figure 3.**
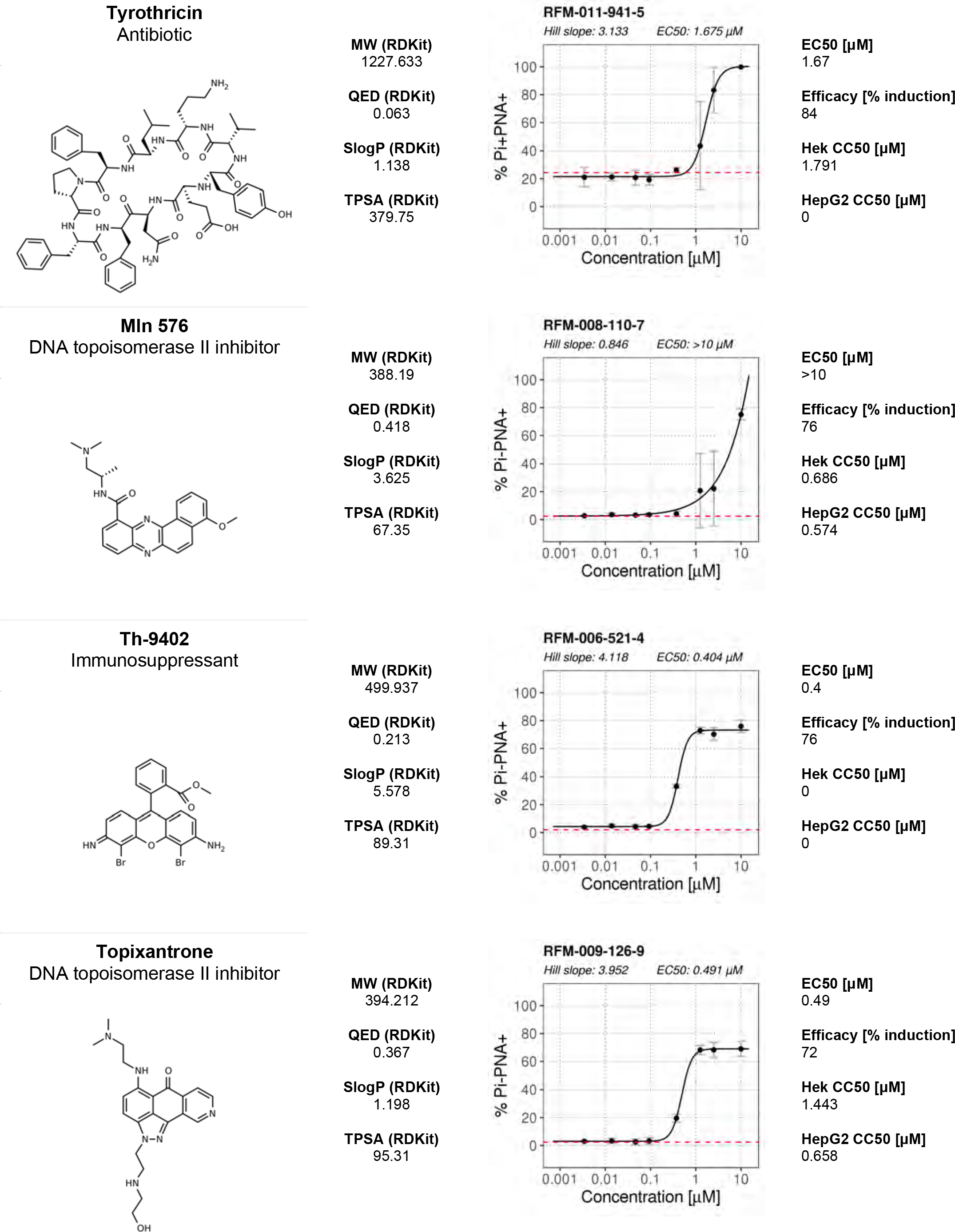
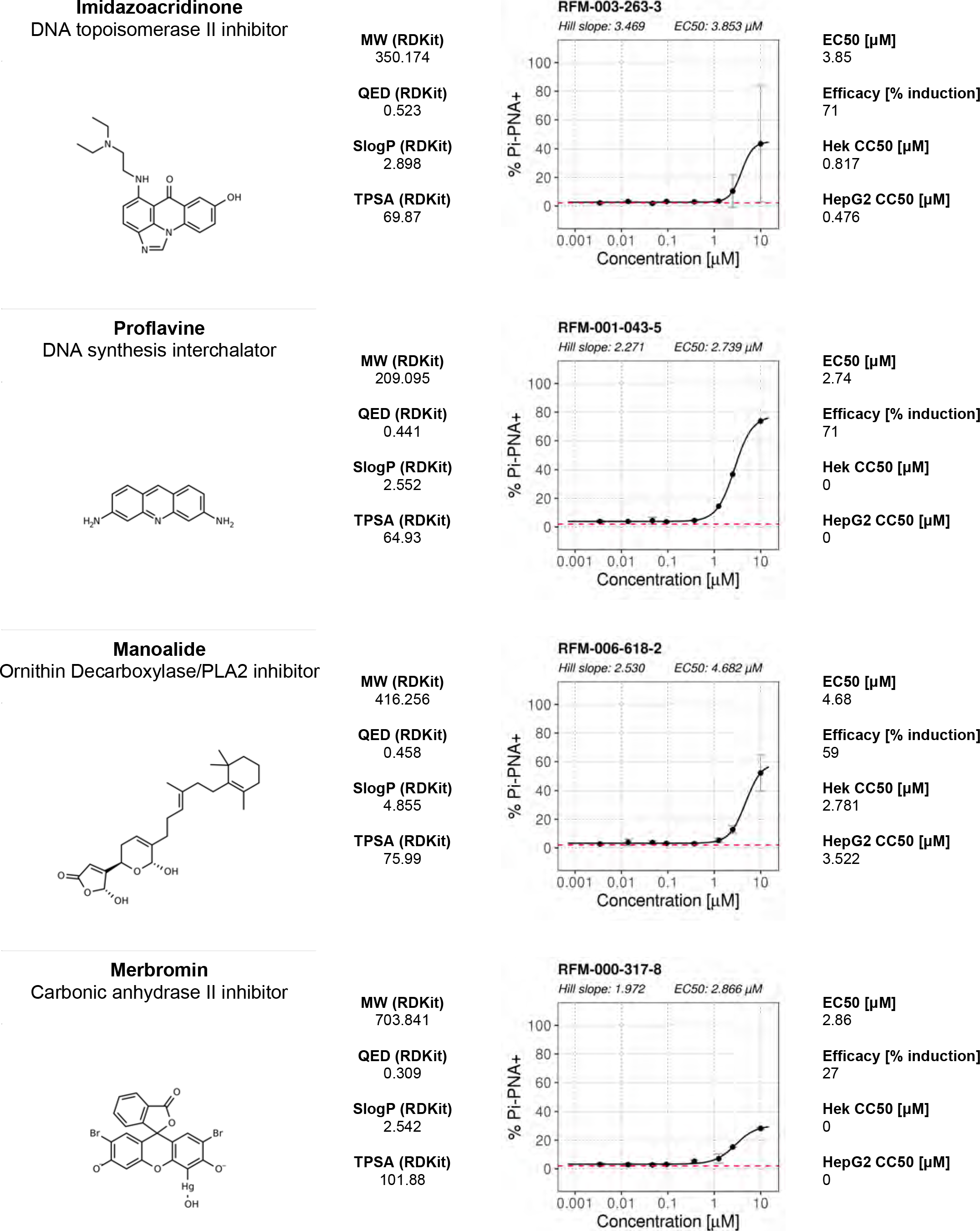
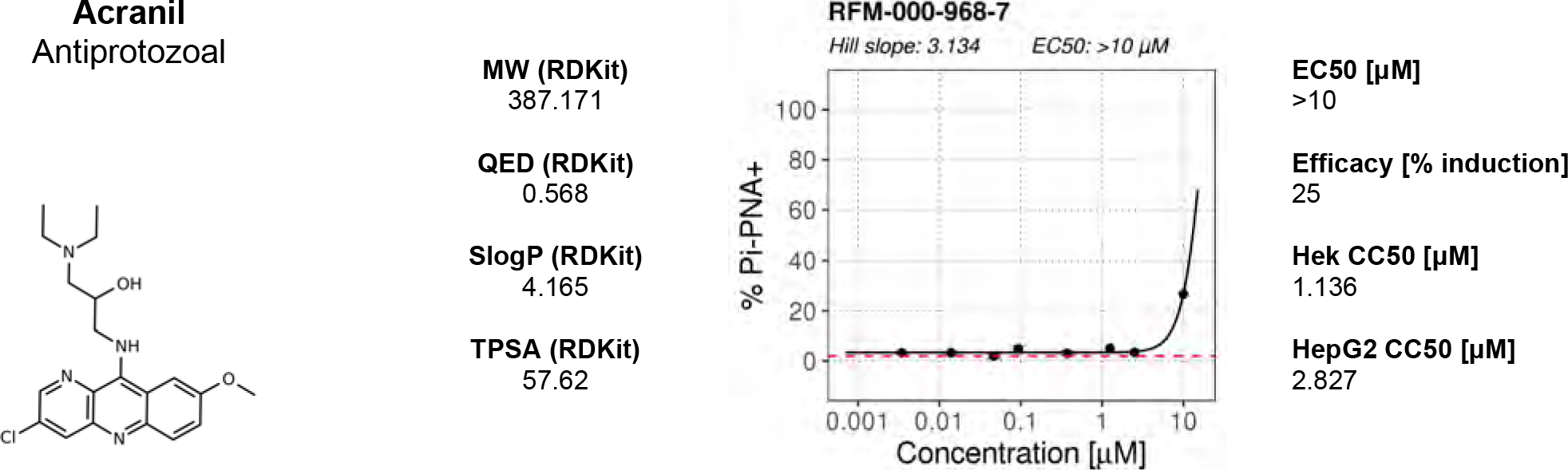
Dose response and additional data for primary AR hits. Compound names and nominal target/functional annotation shown on left along with chemical structure and physico-chemical properties. The 8-point dose response curves for each hit with estimated Hill slope, EC_50_ and Efficacy [% max induction] values are shown on the right. Two data points per concentration (n=2); data points are Mean ± SD. A 4-parameter logistic fit has been performed using R package: *dr4pl*. CC50 values for HEK293 and HepG2 CellTiter-Glo cytotoxicity assays performed by CALIBR (www.reframedb.org) are shown in the right hand section. Note: 0 = inactive in cytotoxicity assay. Physicochemical properties were calculated using RDKit, Python and KNIME: SlogP = partition coefficient (Wildman and Crippen, 1999); TPSA is the Topological Polar Surface Area (Ertl *et al.*, 2000); MW is the exact Molecular weight; QED= Quantitative Estimate of Drug-likeness (Bickerton *et al* 2012). See Supplementary_Figure_3_Source_Data.csv along with Supplementary_Figure_3_Code.R.

**Supplementary Figure 3.1.**
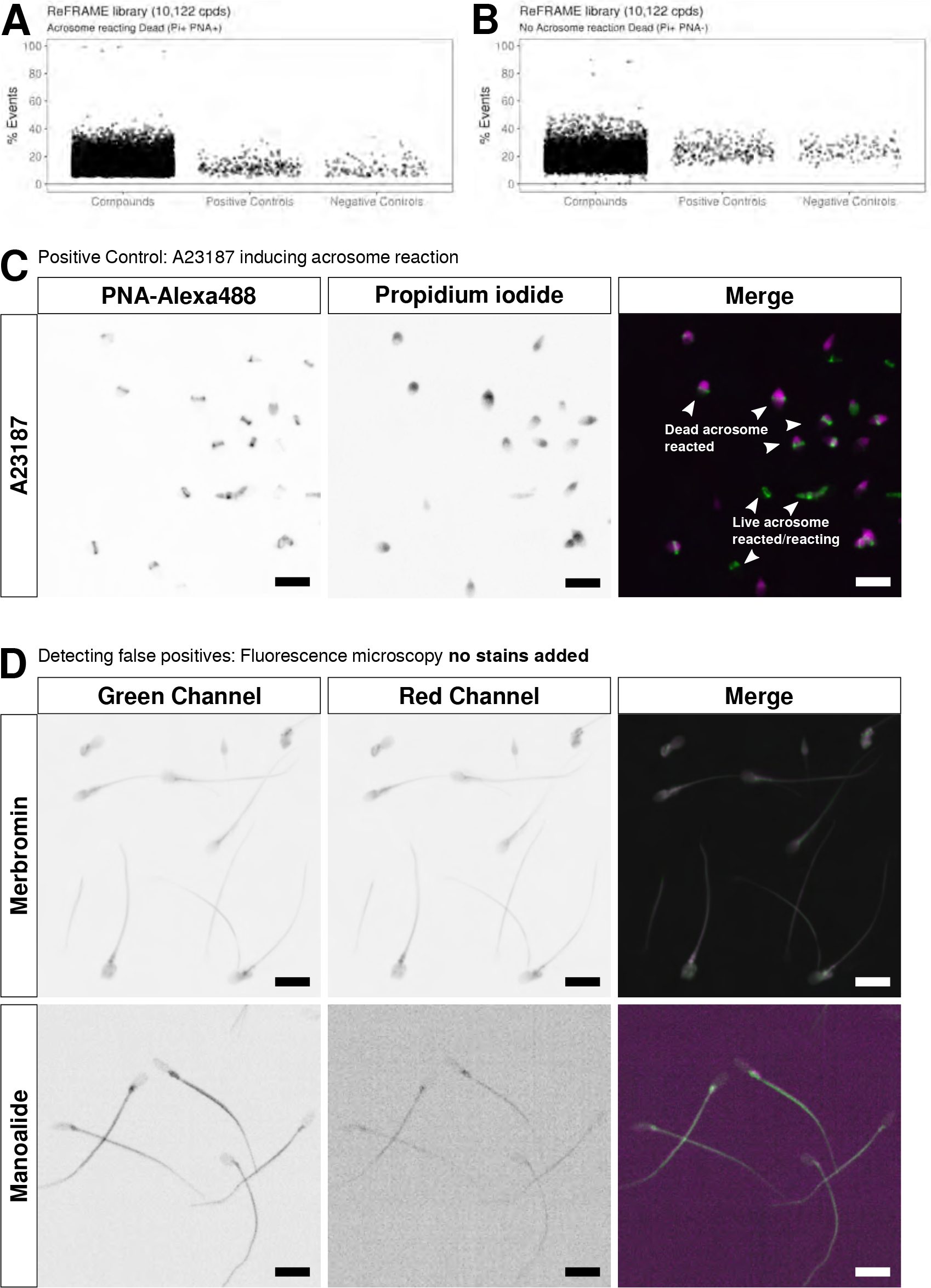
Further analysis of AR screening data and triaging strategy. Results are shown of primary screening data for two of the other populations in the flow cytometry assay: (**A**) Acrosome reaction positive and propidium iodide positive (PI+ PNA+) events and (**B**) acrosome reaction negative and propidium positive (Pi+ PNA-) events. Shown is % Events (number of events relative to total events per well). Datapoints for compounds, negative controls (DMSO) and positive controls (A23187) are shown. See Figure_3_Source_Data.csv along with Figure_3_Code.R. (**C**) Control experiment showing microscopy images of A23187-induced acrosome-reacted sperm. Shown are individual channels (greyscale) for acrosome signal stained with PNA-Alexa488 (green band in merged image) and DNA stained with Propidium iodide indicating lack of cell viability (purple in merged image). Scale bar = 10 μM. (**D**) Orthogonal assay to eliminate fluorescent compounds: Two primary hits were analyzed using fluorescence microscopy without addition of staining reagents: top panels show the non-specific fluorescence of Merbromin and in bottom panels fluorescence of Manoalide which stains only significantly in the midpiece/tail. Grey scale images are shown in the left two panels and a merged image on the right. Key: green channel = Ex 488 nm with Em bandpass filter BP 525/50 nm; red channel = Ex 561 nm; Em Bandpass filter BP600/37 nm). Scale bar = 10 μM.

## Supplementary Movie 1

Movie was generated using a brightfield image sequence for a typical control well (left pane), a well where a compound reduced motility by 20% (middle pane) and one where it reduced it by 80% (right pane).

## Supplementary Movie 2

Movie was generated using a brightfield image sequence for a typical control well (left pane), a well where a compound reduced motility by 20% (middle pane) and one where it reduced it by 80% (right pane). Tracking is overlayed in each panel. Colour coding is detailed in Figure 2.

---

**Supplementary Table 1:**
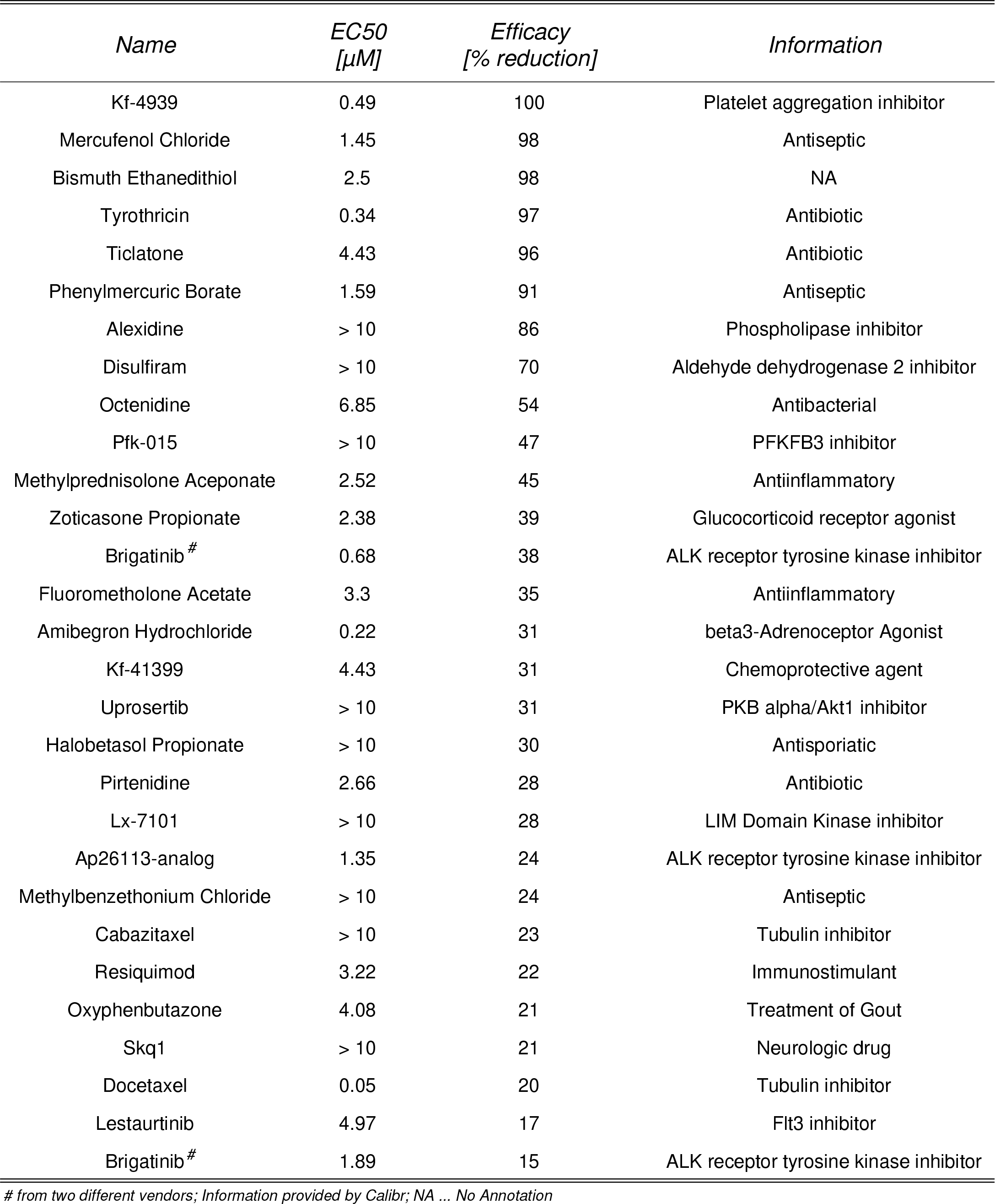
Compounds that had a significant effect on sperm motility. Summary of dose response experiments of primary motility hits with estimated EC50 and Efficacy [% reduction] values. Information and names were provided by Calibr. See Supplementary_Table_1_Source_Data.csv.

| *Name* | *EC50*<br>*[μM]* | *Efficacy*<br>*[% reduction]* | *Information* |
| --- | --- | --- | --- |
| Kf-4939 | 0.49 | 100 | Platelet aggregation inhibitor |
| Mercufenol Chloride | 1.45 | 98 | Antiseptic |
| Bismuth Ethanedithiol | 2.5 | 98 | NA |
| Tyrothricin | 0.34 | 97 | Antibiotic |
| Ticlatone | 4.43 | 96 | Antibiotic |
| Phenylmercuric Borate | 1.59 | 91 | Antiseptic |
| Alexidine | > 10 | 86 | Phospholipase inhibitor |
| Disulfiram | > 10 | 70 | Aldehyde dehydrogenase 2 inhibitor |
| Octenidine | 6.85 | 54 | Antibacterial |
| Pfk-015 | > 10 | 47 | PFKFB3 inhibitor |
| Methylprednisolone Aceponate | 2.52 | 45 | Antiinflammatory |
| Zoticasone Propionate | 2.38 | 39 | Glucocorticoid receptor agonist |
| Brigatinib<sup>#</sup> | 0.68 | 38 | ALK receptor tyrosine kinase inhibitor |
| Fluorometholone Acetate | 3.3 | 35 | Antiinflammatory |
| Amibegron Hydrochloride | 0.22 | 31 | beta3-Adrenoceptor Agonist |
| Kf-41399 | 4.43 | 31 | Chemoprotective agent |
| Uprosertib | > 10 | 31 | PKB alpha/Akt1 inhibitor |
| Halobetasol Propionate | > 10 | 30 | Antisporiatic |
| Pirtenidine | 2.66 | 28 | Antibiotic |
| Lx-7101 | > 10 | 28 | LIM Domain Kinase inhibitor |
| Ap26113-analog | 1.35 | 24 | ALK receptor tyrosine kinase inhibitor |
| Methylbenzethonium Chloride | > 10 | 24 | Antiseptic |
| Cabazitaxel | > 10 | 23 | Tubulin inhibitor |
| Resiquimod | 3.22 | 22 | Immunostimulant |
| Oxyphenbutazone | 4.08 | 21 | Treatment of Gout |
| Skq1 | > 10 | 21 | Neurologic drug |
| Docetaxel | 0.05 | 20 | Tubulin inhibitor |
| Lestaurtinib | 4.97 | 17 | Flt3 inhibitor |
| Brigatinib<sup>#</sup> | 1.89 | 15 | ALK receptor tyrosine kinase inhibitor |

<sup>#</sup> from two different vendors; Information provided by Calibr; NA ... No Annotation

**Supplementary Table 2:** Compounds that had a significant effect on Acrosome Reaction. Summary of dose response experiments of primary acrosome hits with estimated EC50 and Efficacy [% increase] values. Information and names were provided by Calibr. Note that none of these compounds passed orthogonal counter screening and are considered as assay interfering compounds/false positives. See Supplementary_Table_2_Source_Data.csv.

| *Name* | *EC50*<br>*[μM]* | *Efficacy*<br>*[% induction]* | *Information* |
| --- | --- | --- | --- |
| Tyrothricin | 1.67 | 84 | Antibiotic |
| Th-9402 | 0.4 | 76 | Immunosuppressant |
| MIn 576 | >10 | 76 | DNA topoisomerase II inhibitor |
| Topixantrone | 0.49 | 72 | DNA topoisomerase II inhibitor |
| Proflavine | 2.74 | 71 | DNA synthesis interchalator |
| Imidazoacridinone | 3.85 | 71 | DNA topoisomerase II inhibitor |
| Manoalide | 4.68 | 59 | Ornithin Decarboxylase/PLA2 inhibitor |
| Merbromin | 2.86 | 27 | Carbonic anhydrase II inhibitor |
| Acranil | 21.61 | 25 | Antiprotozoal |

Information provided by Calibr

## Source Data Files

*Figure_2_Source_Data.H5*

https://datadryad.org/stash/share/06d75FZ6GiPmme3HnKnkyTFbgKJ2mV0UVRaN-gVKoVE. Primary screening data motility assay

*Figure_3_Source_Data.csv*

Primary screening data acrosome assay

*Supplementary_Figure_2_Source_Data.csv*

Dose response confirmation data motility assay

*Supplementary_Figure_3_Source_Data.csv*

Dose response confirmation data acrosome assay

*Supplementary_Table_1_Source_Data.csv*

Data of Supplementary Table 1

*Supplementary_Table_2_Source_Data.csv*

Data of Supplementary Table 2

## Supplementary Files

*Figure_2_Code.R*

R Code for Figure 2 primary motility assay

*Figure_3_Code.R*

R Code for Figure 2 primary acrosome assay

*Supplementary_Figure_2_Code.R*

R Code for Supplementary Figure 2 dose response confirmation motility assay

*Supplementary_Figure_3_Code.R*

R Code for Supplementary Figure 3 dose response confirmation acrosome assay

## Authors’ roles

FG performed the sperm preparation, HTS screening and analysis of the data. CLRB and PDA designed the study, assisted with interpretation of the data and obtained funding. All authors contributed to the construction, writing, and analysis of data. All authors approved the final manuscript.

## Acknowledgements

We are very grateful to all members of the research team for their invaluable assistance. We also want to thank all the sperm donors who took part in this study and Dr Stephen Gellatly and Ms Morven Dean for recruitment. We acknowledge the assistance of Irene Sucquart in the early phase of this study. Additionally, we want to thank Dr David Mortimer and Dr Sharon Mortimer for their helpful insights into comparisons with the CASA system and advice in AR assay development. Thanks go to Dr Steve Publicover and Dr Zoe Johnston for critical reading of the manuscript. Thanks are also due to NPSC lab members for help, particularly John Raynor for engineering support. We thank Mitch Hull, Emily Chen and Kelli Kunen at CALIBR for their help in library plating, logistics and supply of ReFRAME data. Thanks also go to Professor Kevin Reid, Professor Ian Gilbert and Dr Caroline Wilson of Drug Discovery Unit in Dundee for helpful discussions. Special thanks are due to Professor Andrew Hopkins for his support.

## Funding

Funding was provided by Bill and Melinda Gates Foundation (to PDA and CLRB). CLRB also receives funding from Chief Scientists Office (Scotland) and Astra Zeneca PLC. NPSC was established with funding from the Scottish Funding Council and Scottish Universities Life Science Alliance. PDA/NPSC receives funding from Janssen Pharmaceutica NV.

## Conflict of interest

CLRB is Editor for RBMO, has received lecturing fees from Merck, Pharmasure and Ferring and is on the Scientific Advisory Panel for Ohana BioSciences. PDA, FG had no COI.

